# Fission yeast histone chaperone Rtt106 regulates histone levels, prevents early division, and promotes genome stability

**DOI:** 10.1101/2025.11.01.686071

**Authors:** Samantha A Sanayhie, Victoria Da Pra, Nairy Khodabshakian, Essam Karam, Mohammed Ayan Chhipa, Seerat Kang, Asiya Ahmed, Jeffrey Fillingham, Sarah A Sabatinos

## Abstract

Chromatin packaging influences gene expression and is linked to genome stability through the establishment and maintenance of histone modifications. Histone chaperone proteins regulate chromatin assembly and thus packaging. We tested how loss of the histone chaperone Rtt106 affects genome stability through cell cycle checkpoint stability in response to cellular stress. We tested how double mutants lacking DNA replication or DNA damage checkpoint kinases are impacted by the absence of histone chaperone *rtt106*. Rtt106 brings histone H3 and histone H4 together into complexes. We found that *rtt106*Δ cells with loss of the DNA replication checkpoint (*cds1*Δ*, rad3*Δ) were more sensitive to hydroxyurea. However, DNA damage kinase *chk1*Δ *rtt106*Δ cells became less sensitive to DNA damaging drugs. The effects of Rtt106 on growth are observed in division timing, where *rtt106*Δ cells show early division in the presence of drug. Coupled to a decrease in histone H3 levels and increased mutation rate, our work shows how non-essential Rtt106 activities contribute to genome stability. By regulating histone levels and use, Rtt106 regulates the cell division and may function in chromatin arm coherence and segregation.

## INTRODUCTION

Chromatin formation and folding are two processes that regulate DNA transcription and impact genome stability (Felsenfeld, 1978; Gross *et al*., 2015). The ability to form chromatin by winding DNA around a protein complex allows the entire genome to fit within a nucleus. Chromatin prevents DNA damage, while restricting access for enzymes that may contribute to genome instability (Kouzarides, 2007; Campos and Reinberg, 2009; Gross *et al*., 2015). Thus, chromatin maintenance is important because DNA damage and mutation can lead to altered cell growth, improper chromosome segregation, cellular aging, and diseases such as cancer.

Histone chaperones direct the formation and maintenance of chromatin. The basic subunits of chromatin are nucleosomes, which are units of 147 DNA bp encircling a histone core (Kornberg, 1974; Gross *et al*., 2015). The histone core is an octamer, containing two histone H2A-H2B dimers and one histone H3-H4 tetramer. The formation of the H2A-H2B and (H3-H4)_2_ subunits and then assembly of the octamer complex are assisted by specific histone chaperones. Chaperones also mediate DNA-octamer association and assembly (Fazly *et al*., 2012; Mattiroli *et al*., 2015; Hammond *et al*., 2017). Chromatin assembly steps are regulated by chaperone protein structural folds and domains, promoting proper histone deposition and histone formation (Burgess and Zhang, 2013; Gurard-Levin *et al*., 2014; Mattiroli *et al*., 2015; Hammond *et al*., 2017).

Histone chaperone Rtt106 binds to the (H3-H4)_2_ tetramer and directs its deposition into the final octamer form, and subsequently onto the DNA (Huang *et al*., 2005; Fazly *et al*., 2012). Rtt106 can oligomerize and stabilize (H3-H4)_2_ tetrameric interactions protecting the DNA from damage (Fazly *et al*., 2012; Su *et al*., 2012). Because of these interactions between Rtt106, DNA, and damage maintenance, we predicted that loss of Rtt106 may promote genome instability. To test this hypothesis, we generated a *rtt106*Δ strain in *Schizosaccharomyces pombe.* We then tested how the loss of Rtt106 affects cell health and fitness in cell cycle checkpoint mutants.

We tested checkpoint kinase mutants to assess interactions between Rtt106 function and DNA damage and replication stress. Checkpoint kinases respond to DNA damage and genome instability. In fission yeast, the human ATR-homologue Rad3 is activated in response to DNA replication stress or DNA damage (McCULLY and Robinow, 1971; Lindsay *et al*., 1998; Martinho *et al*., 1998; Caspari and Carr, 1999; Sabatinos *et al*., 2012; Willet *et al*., 2015). The intra-S and S/M checkpoints ensure DNA is appropriately and correctly replicated (Lindsay *et al*., 1998; Martinho *et al*., 1998; Sabatinos and Forsburg, 2010; Barnum and O’Connell, 2014). During replication instability, single stranded DNA accumulation slows DNA replication forks and targets Rad3 to phosphorylate the Cds1 kinase. Cds1 is the main replication stress effector kinase that arrests the cell cycle by Cdc2 phosphorylation, blocks DNA origin firing, and promotes stability (Forsburg and Nurse, 1991; Martinho *et al*., 1998; Caspari and Carr, 1999; Barnum and O’Connell, 2014) that allow later repair and replication restart. Removing either Rad3 or Cds1 (*cds1*Δ*, rad3*Δ strains) makes cells sensitive to DNA replication stress. The *cds1*Δ or *rad3*Δ cells cannot survive when hydroxyurea (HU) causes dNTP depletion and fork slowing or arrest. We predicted that *cds1*Δ *rtt106*Δ or *rad3*Δ *rtt106*Δ cells would be more sensitive to HU.

The fission yeast G2/M checkpoint is activated if DNA damage is present prior to mitosis. DNA damage can be induced *in vitro* by camptothecin (CPT) or phleomycin (phleo). After DNA breaks are resected, the resulting ssDNA activates the Rad3 kinase. Rad3 phosphorylates the Chk1 effector kinase, which phosphorylates Cdc2 and activates repair pathways (Martinho *et al*., 1998; Caspari and Carr, 1999; Kastan and Bartek, 2004; Sabatinos *et al*., 2012; Barnum and O’Connell, 2014). Removing either Rad3 or Chk1 (*chk1*Δ*, rad3*Δ strains) makes cells sensitive to DNA damage. The *chk1*Δ or *rad3*Δ cells cannot survive double strand breaks generated in CPT or phleo. We predicted that *chk1*Δ *rtt106*Δ or *rad3*Δ *rtt106*Δ cells would be more sensitive to DNA damage caused by drug because of altered chromatin stability.

We report that Rtt106 loss increases histone H3 levels, leading to increased mutation rates. Drug sensitivity, cell growth and cell division rates show that Rtt106 functions impact DNA replication and repair checkpoint outcomes. Importantly, *rtt106*Δ cells show clear differences in cell division timing and frequency. Our data suggests that an overall increase in histone H3 levels of *rtt106*Δ contribute to DNA damage in an otherwise checkpoint competent background. However, the loss of the DNA repair checkpoint causes precocious cell division. We conclude that the increased DNA damage in *rtt106*Δ promote increased mutagenesis. Together, we show that Rtt106 plays a role in maintaining genomic stability.

## RESULTS

*S. cerevisiae* RTT106 binds to the HIR complex and generates repressive chromatin that stops transcription of core histones. Thus, histone HTA1 RNA levels are reduced in the *S. cerevisiae rtt106*Δ mutant (*Sc rtt106*Δ), causing altered nucleosome occupancy at HIR-responsive promoters (Fillingham *et al*., 2009). Because *Sc* RTT106 is different from *S. pombe* Rtt106 in both regulation and potential function, we made a *rtt106*Δ strain by targeted integration of the KanMX6 resistance gene into *rtt106^+^.* Our construct deleted the protein coding region of *rtt106^+^* (Supplementary Figure S1). Integration was confirmed by PCR to show that *rtt106* coding sequence was lost upon KanMX6 integration into *rtt106*-flanking regions. To test whether *S. pombe* Rtt106 influences histone levels and genome stability (Figure 1A), we assessed histone H3 level by western blot compared to a PCNA loading control (Figure 1B). Increased histone H3 in *rtt106*Δ cells confirms that Rtt106 function impacts overall histone level and may impact genome stability in *S. pombe*.

**Figure 1.**
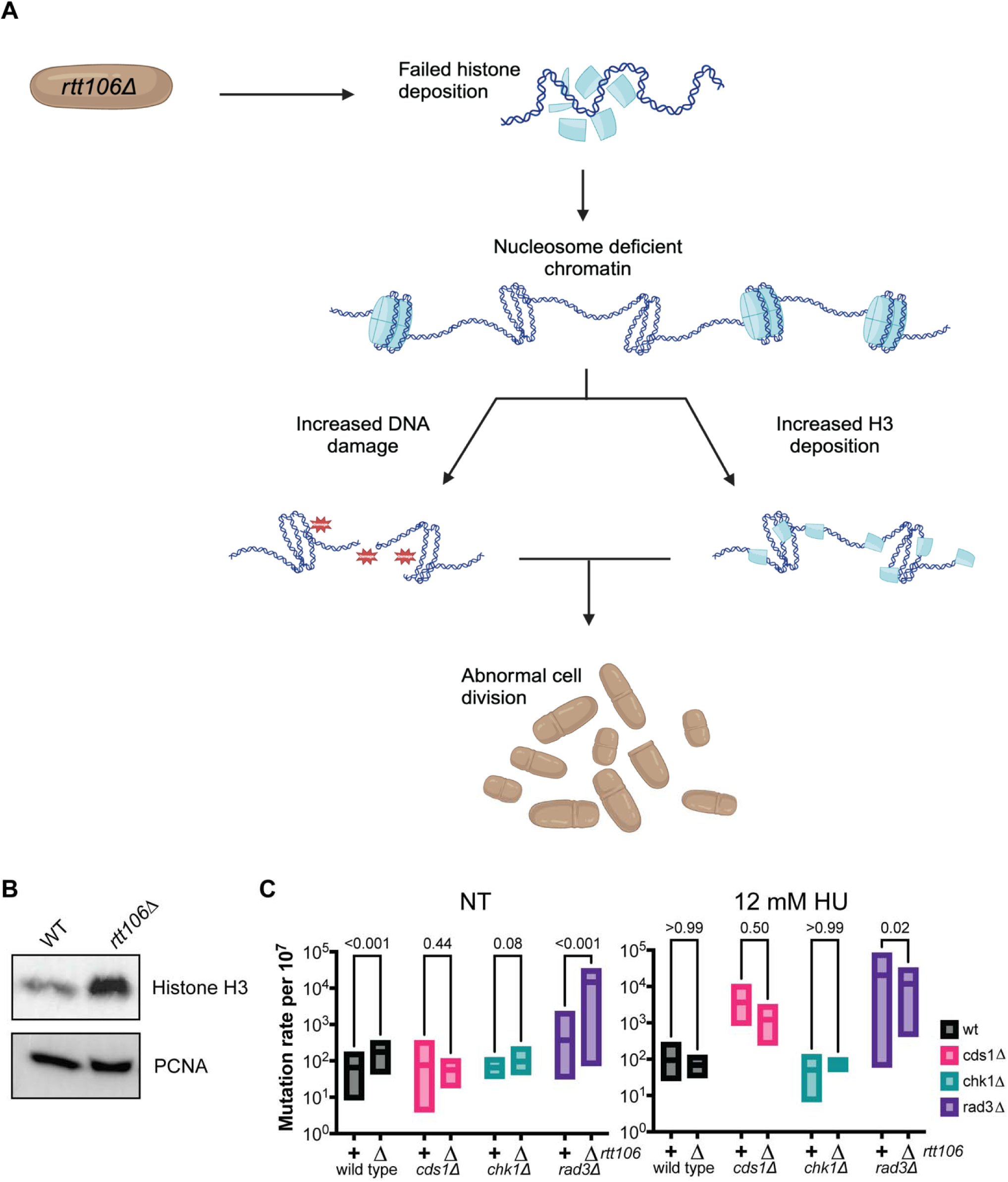
A role for *S. pombe* Rtt106 in genome stability. A) In addition to Rtt106 roles in H3-H4 dimer and tetramer formation, *S. cerevisiae* Rtt106 directs nucleosome deposition and represses chromatin at core histones. We hypothesized that *S. pombe* Rtt106 similarly affects histone level. We predict that *rtt106*Δ cells may have altered histone level and/or increased DNA damage that results in abnormal mitoses and genome instability. B) Western blot of histone H3 protein levels were measured in wild type (WT) and *rtt106*Δ strains, with PCNA as a control. Increased H3 was seen in asynchronous cultures of *rtt106*Δ cells compared to WT. C) Canavanine forward mutation rate in untreated (left) or HU-treated (right) cultures. The Lea and Coulson mutation rate per 10^7^ generations is shown with box of entire data set (n=5+ independent replicates). Refer to Table 2 for MSS-MLE mutation rate estimate and significance results.

**Table 1:** Yeast Strains used in this study.

| Name (strain number) | Genotype | Source |
| --- | --- | --- |
| Wild type (SY17) | *h<sup>+</sup> leu1-32::hENT-leu1<sup>+</sup> his7-366::hsv-tk-his7<sup>+</sup> ura4-D18 ade6-M21?* | S Forsburg (FY2317) |
| *cds1Δ* (SY91) | *h<sup>+</sup> rtt106Δ::kanMX6 cds1Δ::ura4<sup>+</sup> leu1-32::hENT-leu1<sup>+</sup> his7-366::hsv-tk-his7<sup>+</sup> ura4-D18 ade6-M21?* | S Forsburg (FY5148) |
| *chk1Δ* (SY92) | *h<sup>+</sup> rtt106Δ::kanMX6 chk1Δ::ura4<sup>+</sup> leu1-32::hENT-leu1<sup>+</sup> his7-366::hsv-tk-his7<sup>+</sup> ura4-D18 ade6-M21?* | S Forsburg (FY5149) |
| *rad3Δ* (SY93) | *h<sup>+</sup> rtt106Δ::kanMX6 rad3Δ::ura4<sup>+</sup> leu1-32::hENT-leu1<sup>+</sup> his7-366::hsv-tk-his7<sup>+</sup> ura4-D18 ade6-M21?* | S Forsburg (FY5150) |
| *rtt106Δ* (SY153) | *h<sup>-</sup> rtt106Δ::KanMX ade6-M21? leu1-32 his6-366 ura4-D18 ade6-M216* | this study |
| *rtt106Δ cds1Δ* (SY268) | *h<sup>+</sup> rtt106Δ::kanMX6 cds1Δ::ura4<sup>+</sup> leu1-32::hENT-leu1<sup>+</sup> his7-366::hsv-tk-his7<sup>+</sup> ura4-D18 ade6-M21?* | this study |
| *rtt106Δ chk1Δ* (SY153) | *h<sup>+</sup> rtt106Δ::kanMX6 chk1Δ::ura4<sup>+</sup> leu1-32::hENT-leu1<sup>+</sup> his7-366::hsv-tk-his7<sup>+</sup> ura4-D18 ade6-M21?* | this study |
| *rtt106Δ rad3Δ* (SY153) | *h<sup>+</sup> rtt106Δ::kanMX6 rad3Δ::ura4<sup>+</sup> leu1-32::hENT-leu1<sup>+</sup> his7-366::hsv-tk-his7<sup>+</sup> ura4-D18 ade6-M21?* | this study |

**Table 2:** Canavanine mutation rate in cells with and without 12 mM HU.

|  | Untreated MSS-MLE mutation rate per 10<sup>7</sup> generations |  | MSS-MLE mutation rate per 10<sup>7</sup> generations, after 12 mM HU, 4h, 30°C |  |
| --- | --- | --- | --- | --- |
|  | *rtt106<sup>+</sup>* | *rtt106Δ* | *rtt106<sup>+</sup>* | *rtt106Δ* |
| Wild type | 135 | 162 | 92.4 | 67.2 |
| *cds1Δ* | 41.0 | 49.8 | 2542 | 841 |
| *chk1Δ* | 68.6 | 81.9 | 33.0 | 77.7 |
| *rad3Δ* | 85.0 | 6446 | 8452 | 11256 |

DNA mutagenesis is often increased in conditions of genome instability, and we hypothesized that *rtt106*Δ cells may show increased mutation rates. We plated onto canavanine, which is a toxic metabolite that kills cells that have functional amino acid transport of arginine and lysine (Rosenthal, 1977, 2001). Mutations that alter transport may allow cells to grow and form colonies on canavanine plates (Ait Saada *et al*., 2022; Yang *et al*., 2022; Lyu *et al*., 2024). We found that *rtt106*Δ mutation rates were significantly higher than *rtt106^+^* wild type cultures (Figure 1C, Table 2).

We asked whether cell cycle checkpoints regulate DNA damage or replication instability in *rtt106*Δ cells that causes mutation. By crossing the *rtt106*Δ allele into checkpoint mutants (*rad3*Δ*, chk1*Δ*, cds1*Δ), we found that the canavanine mutation rate was almost 100-times higher in *rad3*Δ *rtt106*Δ cells compared to *rad3*Δ alone. The *chk1*Δ and *chk1*Δ *rtt106*Δ cells may also show mildly higher mutation rates (p∼0.08). These data imply that Rtt106 plays roles in some mechanisms of genome stability.

Because increased mutation rates could reflect a role of Rtt106 in managing DNA replication, we added 12 mM hydroxyurea (HU) to growing cells for 4h, depleting dNTPs and inducing replication instability (Figure 1C). The presence of Rtt106 had no significant effect on canavanine mutation rate in wild type, *cds1*Δ, or *chk1*Δ cells. HU-exposure mutation rate was higher in *cds1*Δ and *cds1*Δ *rtt106*Δ cells compared to no HU. Yet, *rad3*Δ mutation rate in HU was higher *rtt106+* cells. The *rad3*Δ *rtt106*Δ mutation rate was similar with or without HU. Thus, the effect of HU on mutation rate is epistatic to *rtt106*Δ alone, and there is no increase in *rad3*Δ *rtt106*Δ mutation specifically caused by HU.

### Drug survival is increased in some rtt106Δ mutant cells

Because our mutation rate data suggested that *rtt106*Δ enhances genome instability without drug treatment, we next asked how whether Rtt106 functions in the DNA replication or DNA damage checkpoints. We spotted dilutions of single and double mutant *rtt106*Δ strains on media containing: hydroxyurea (HU) to induce DNA replication stress and S-phase arrest (Musiałek and Rybaczek, 2021); camptothecin (CPT) to cause topoisomerase trapping and DNA breaks in subsequent S-phases (Liu *et al*., 2000); and, phleomycin (phleo) to generate DNA double strand breaks independently of cell cycle stage (Sleigh, 1976). We found that *rtt106*Δ cells generally grew like *rtt106^+^*cells on low-dose and chronic concentrations of the three drugs (Figure 2A).

**Figure 2.**
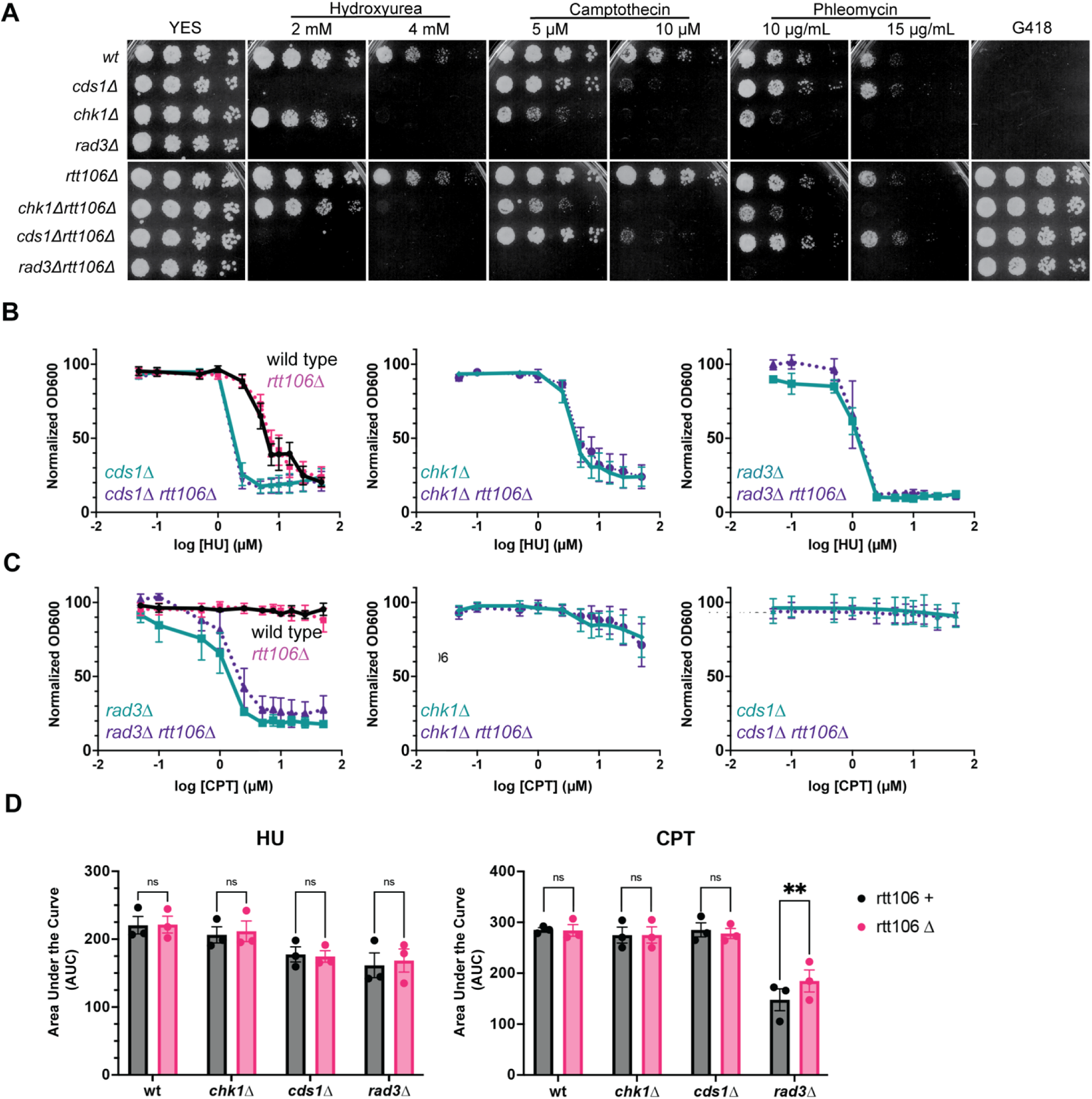
Rtt106 has mild effects on checkpoint mutant response to drug. A) Strains were assessed for drug sensitivities using a spot test on solid YES medium. Strains were normalized at 5 x 10^6^ cells/mL and diluted in four 1:5 dilutions. The *rtt106*Δ grew similarly to wild type (wt) under all drug conditions (hydroxyurea, camptothecin, and phleomycin). Double mutant checkpoint strains with *rtt106*Δ had similar drug sensitivities to their single mutant checkpoint parental strain. B,C) Growth in YES liquid cultures was modeled by monitoring OD600 at a range of hydroxyurea (B) and camptothecin (C) doses. Shown are dose-response curves for each strain. A minimum of 3 experimental replicates were normalized to respective no drug value, and then modeled with a log(inhibitor) response and variable slope. Shown is the composite response across replicate data for each strain and condition. In HU, matched *rtt106*Δ strains behaved like the corresponding *rtt106+* strain. However, variation in *rad3*Δ *rtt106*Δ versus *rad3*Δ indicated that subtle changes occur in HU that are not captured by OD600 liquid cultures. C) IC50 curves in CPT are similar in wild type, *cds1*Δ, and *chk1*Δ pairs (*rtt106^+^ vs rtt106*Δ). The *rad3*Δ *rtt106*Δ cells grow better in CPT than *rad3*Δ alone (p=0.016). IC50 values are presented in Table XX. D) Area under the curve (AUC) was used to model differences in replicate growth curves. HU decreases growth in all matched *rtt106+ vs rtt106*Δ pairs. CPT effects on growth curves are not significant in wild type, *cds1*Δ, and *chk1*Δ. An exception is that *rad3*Δ *rtt106^+^* cells are more sensitive to CPT than *rad3*Δ *rtt106*Δ, which has a significant impact in AUC (** p<0.01).

Effects were subtle. For example, *cds1*Δ *rtt106*Δ grew slightly better than *cds1*Δ on 10 µM CPT or 15 µg/mL phleo. The *rtt106*Δ also grew better than wild type (*rtt106+)* on 10 µM CPT. Hydroxyurea effects were negligible, and there was no apparent difference between *rad3*Δ and *rad3*Δ *rtt106*Δ. We inferred that *rad3*Δ and *rad3*Δ *rtt106*Δ strains were both sensitive at the doses tested, and that Rtt106 has a minor role in altering DNA damage sensitivity.

Because small differences in the growth of cells within spots can be misleading, we tested ranges of HU and CPT doses on *rtt106*Δ proliferation. We measured optical density of liquid cultures and calculated then compared the half-maximal inhibitory dose (IC50) for each strain (Figure 2B, 2C). The area under each curve (AUC) was used to assess the impact of genotype on overall growth in HU or CPT (Figure 2D). We found that IC50 doses were similar for *rtt106^+^ and rtt106*Δ pairs in HU (Figures 2B, 2D, Table 3), and the HU AUCs were similar (Figure 2D). The *rad3*Δ *rtt106*Δ cells grew slightly better in HU compared to *rad3*Δ alone.

**Table 3:**
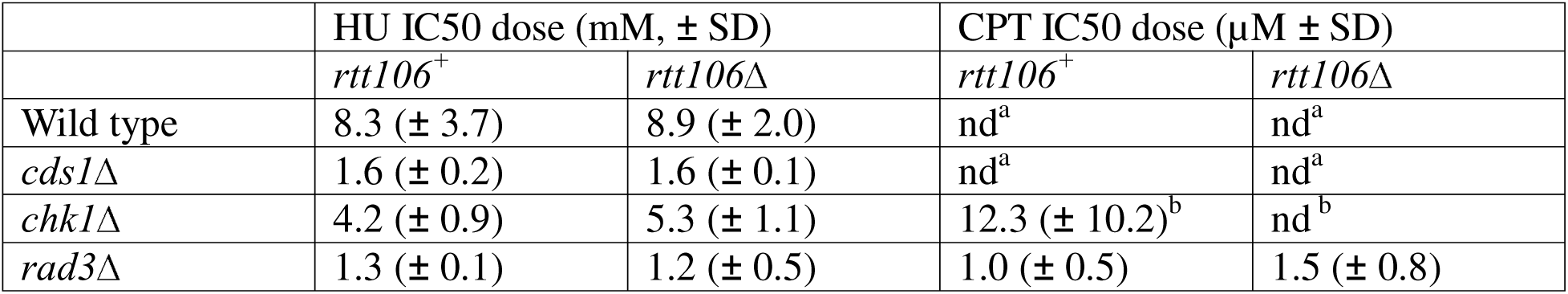
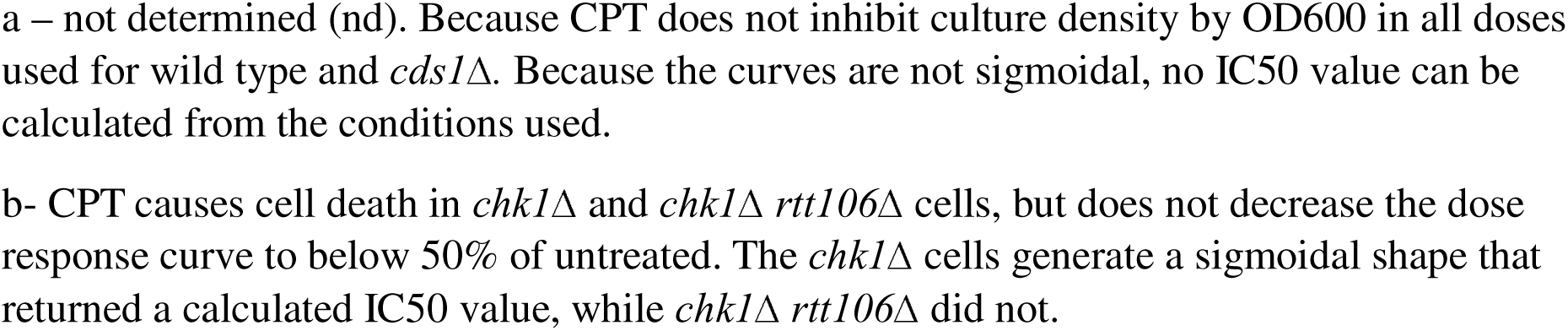
IC50 values for strains in HU and CPT (refer to Figure 1 for averaged curves from at least 3 experimental replicates).

|  | HU IC50 dose (mM, ± SD) |  | CPT IC50 dose (μM ± SD) |  |
| --- | --- | --- | --- | --- |
|  | *rtt106<sup>+</sup>* | *rtt106Δ* | *rtt106<sup>+</sup>* | *rtt106Δ* |
| Wild type | 8.3 (± 3.7) | 8.9 (± 2.0) | nd<sup>a</sup> | nd<sup>a</sup> |
| *cds1Δ* | 1.6 (± 0.2) | 1.6 (± 0.1) | nd<sup>a</sup> | nd<sup>a</sup> |
| *chk1Δ* | 4.2 (± 0.9) | 5.3 (± 1.1) | 12.3 (± 10.2)<sup>b</sup> | nd<sup>b</sup> |
| *rad3Δ* | 1.3 (± 0.1) | 1.2 (± 0.5) | 1.0 (± 0.5) | 1.5 (± 0.8) |

a – not determined (nd). Because CPT does not inhibit culture density by OD600 in all doses used for wild type and *cds1* $\Delta$ . Because the curves are not sigmoidal, no IC50 value can be calculated from the conditions used.
b- CPT causes cell death in *chk1* $\Delta$ and *chk1* $\Delta$ *rtt106* $\Delta$ cells, but does not decrease the dose response curve to below 50% of untreated. The *chk1* $\Delta$ cells generate a sigmoidal shape that returned a calculated IC50 value, while *chk1* $\Delta$ *rtt106* $\Delta$ did not.

In CPT, most strains were not sensitive at the doses we used, and their IC50 is listed as >500 µM (Figure 2C, Table 3). The *rad3*Δ *rtt106*Δ strain grew more in CPT than its *rad3*Δ parental partner, reflected in higher OD600 values and higher AUC (Figure 2C, 2D). The CPT IC50 dose for *rad3*Δ was 1.27 µM, which was lower than the CPT IC50 of *rad3*Δ *rtt106*Δ at 1.45 µM. Lower IC50 shows that *rad3*Δ is more sensitive than *rad3*Δ *rtt106*Δ. While we had not seen a difference in *rad3*Δ CPT sensitivity by spot tests, the IC50 test allowed us to determine changes in growth between *rad3*Δ and *rad3*Δ *rtt106*Δ. Additionally, the *chk1*Δ *rtt106*Δ mutant cells generated a higher IC50 dose of 5.3 mM, compared to *chk1*Δ at 4.2mM HU, underscoring small but intriguing differences between double and single mutant cultures.

### Growth during replication stress is impacted by rtt106 loss

To see whether Rtt106 impacts growth during an acute DNA replication stress response, we tested single and double mutant strains in varying doses of hydroxyurea. In fission yeast, one cell cycle takes approximately 2.5 hours (Carlson *et al*., 1999). DNA replication or DNA damage stress either arrest the cell cycle during drug treatment, or cause cell death if the appropriate checkpoint is absent. We investigated Rtt106 roles in cell growth kinetics, cell length, and cell division rates in the presence of varying hydroxyurea doses (0, 3, 6, 12 mM) over 24 h. Growth curves of all strains show small differences between *rtt106*Δ and matched parental strains (Figure 3A, 3B, 3C). Intriguingly, *chk1*Δ *rtt106*Δ and *rad3*Δ *rtt106*Δ cells show increased OD600 at 12 mM HU, and slightly higher at 6 mM HU. Because fission yeast elongate during HU exposure with accumulation of cells in G2 arrest, the higher OD600 could indicate either faster divisions with more cells or longer cells. We tested for cell length by measuring cells from each genotype before and after 12 mM HU for 6h. We found that wild type cells are significantly longer in HU, but that *rtt106*Δ cells are not (Figure 3D). This may contribute to the slight decrease of *rtt106*Δ OD600 compared to wild type in Figure 3A. Cells were longer in *chk1*Δ*, chk1*Δ *rtt106*Δ*, cds1*Δ, and *cds1*Δ *rtt106*Δ after 6h in 12 mM HU; yet, the patterns with and without *rtt106* were not different. Similarly, both *rad3*Δ and *rad3*Δ *rtt106*Δ cells are smaller after 6h HU.

**Figure 3.**
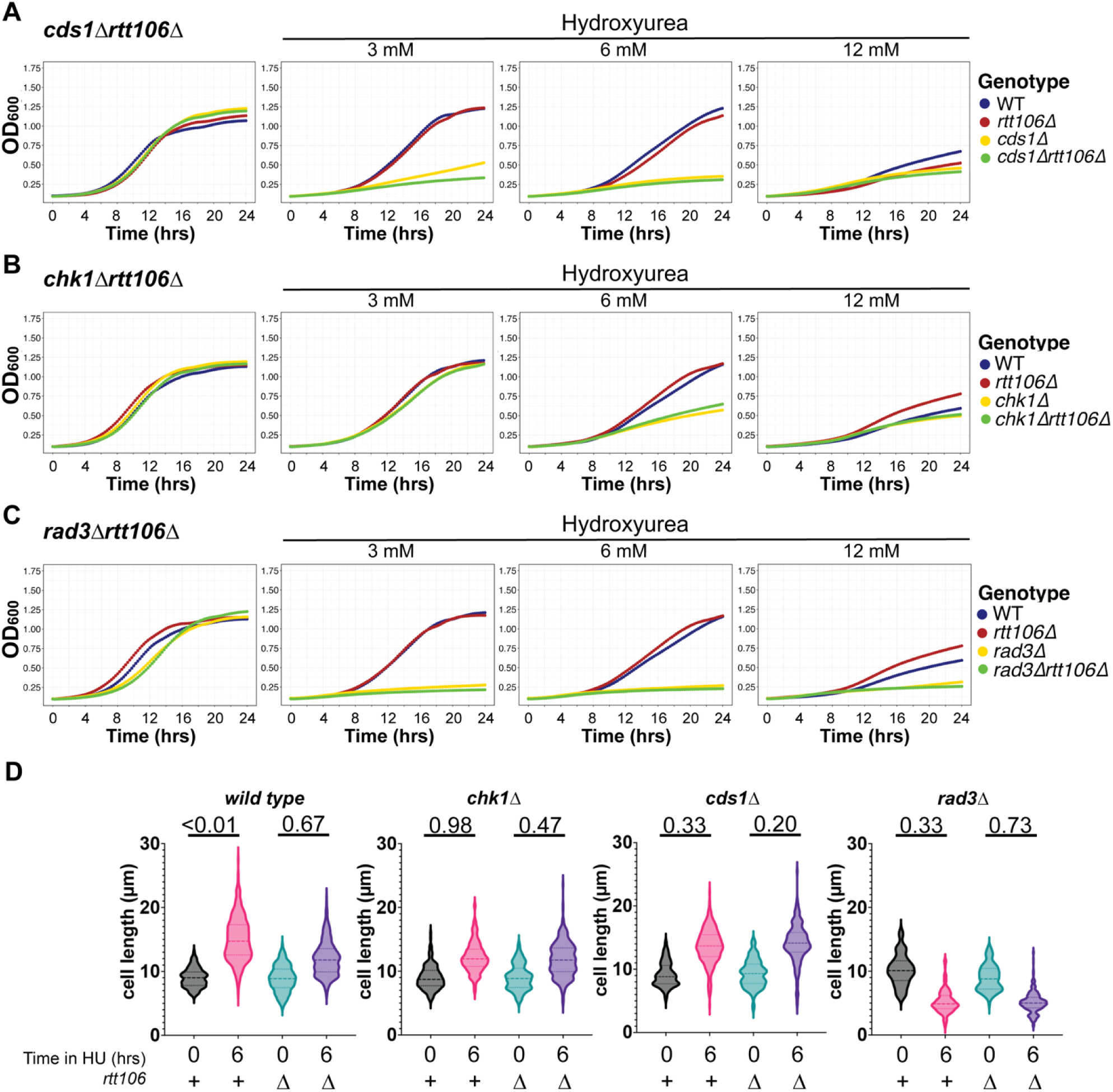
Culture growth during short-term DNA replication stress is similar in *rtt106*Δ single and double mutants. A, B, C) Growth kinetics were assessed in YES containing 3 mM, 6 mM, or 12 mM hydroxyurea, and compared to no HU over 24 hours. OD600 was used to monitor turbitidity as a proxy for concentration. Wild type and *rtt106*Δ cells exhibit very similar growth patterns under all drug concentrations; these are shown in each plot, matched to the experiment of the paired strains. Representative curves showing an example from one of 3+ experimental replicates. Matched pairs of *rtt106+* and *rtt106*Δ cultures in *cds1*Δ (A), *chk1*Δ (B), and *rad3*Δ (C) are compared to checkpoint kinase wild type. We found that *cds1*Δ*, cds1*Δ *rtt106*Δ*, rad3*Δ, and *rad3*Δ *rtt106*Δ cells are all exquisitely sensitive to HU during growth. The *chk1*Δ and *chk1*Δ *rtt106*Δ cells are more sensitive to 6 mM HU than wild type and *rtt106*Δ; at 3 mM and 12 mM HU the *chk1*Δ pair grows like wild type. D) Cell length of cells after 12 mM HU for 6h (YES, 30°C) is compared with the asynchronous 0h cells. Wild type cells elongate in HU (p<0.01), but *rtt106*Δ lengths are not significantly changed after 6h in HU. Other matched pairs did not show significant differences in cell lengths across populations. Shown are violin plots showing the cell length distributions of 75-100 cells sampled in a minimum of 3 experimental replicates (n=3 for statistics).

Because increased mutation rate and elongation in wild type could correlate with changed viability, we tested colony formation in cells treated with 12 mM HU (Figure 4A). Wild type and *chk1*Δ cells have a functional DNA replication checkpoint and should survive acute HU exposure and form colonies. Consistent with spot tests and IC50 data, we found that there was no change in relative viability depending on *rtt106* during acute HU exposure. Replication checkpoint deficient *rad3*Δ and *cds1*Δ cells should show lower acute viability and fewer colonies in HU. The viability of *rad3*Δ and *cds1*Δ did not change with Rtt106 loss (*rtt106*Δ). We concluded that the difference in OD600 growth for *chk1*Δ*/chk1*Δ*rtt106* and *rad3*Δ */ rad3*Δ *rtt106*Δ pairs was independent of cell length and reflects an alternative mechanism that may impact genome stability.

**Figure 4.**
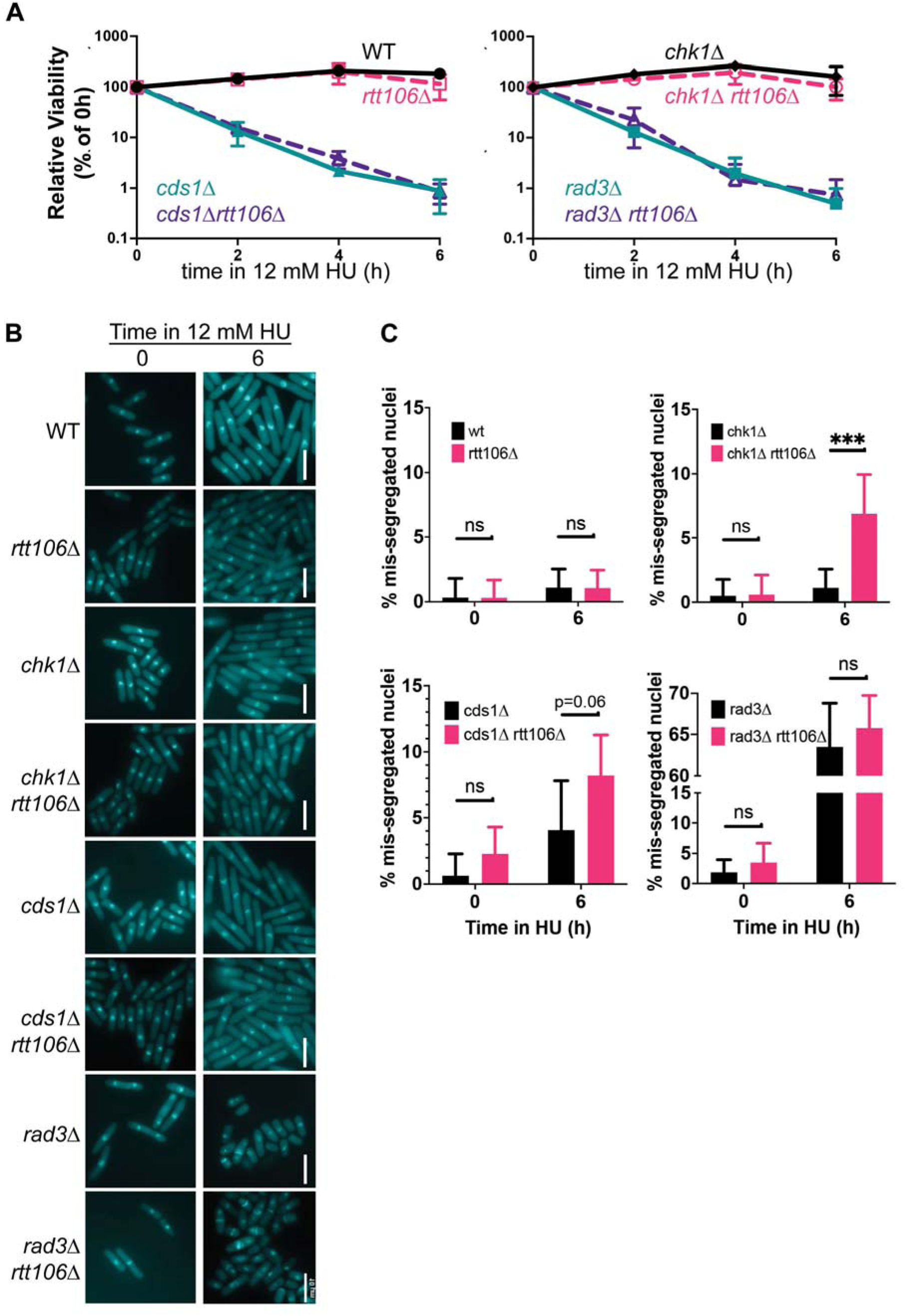
Rtt106 prevents DNA mis-segregation caused by acute hydroxyurea exposure in DNA damage checkpoint mutant *chk1*Δ. A) Acute replication stress exposure (12 mM HU, 30°C) was monitored by plating samples for colony formation over time. Shown is the average relative viability for each strain (n=3). during acute exposure at 30°C. B) Cells were stained with aniline blue and DAPI to detect septa and nuclei, comparing before and after 6 h in 12 mM HU. Scale bars are 10 µm. C) Mis-segregated nuclei were counted as a proportion of cell populations, and compared at no HU, or 6h HU for each genotype. A minimum of 50 cells from at least 3 images were counted over experimental replicates. Mis-segregation was scored for anucleate (no DNA in a cell), lagging DNA, *cut*, or micronuclei. Shown are the proportion of pooled data across experiments with 95% confidence interval. The percent of mis-segregation was compared between paired genotypes using a chi-square test.

To understand gross genome stability effect of *rtt106*, we assessed DNA segregation before and after 12 mM HU using DAPI staining for DNA, and aniline blue for septa (Figure 4B). We counted the number of DNA mis-segregation events in populations of cells including lagging chromosomes, micronuclei, cell untimely torn “*cut*”, and anucleate cells (Figure 4C). We found that there was a significantly higher proportion of mis-segregated nuclei in *chk1*Δ *rtt106*Δ cells. Other differences were not below p<0.05. For example, *rad3*Δ *rtt106*Δ cells had similarly mis-segregated nuclei compared to *rad3*Δ. The *cds1*Δ *rtt106*Δ cells had higher mis-segregation, although the p-value was 0.06. We concluded that abnormal DNA division may contribute to the difference in 24h HU growth of *chk1*Δ *rtt106*Δ and *rad3*Δ *rtt106*Δ (Figure 3B, 3C).

### Rtt106 stops mitosis during replication stress or DNA damage

To test how mutation might occur in *rad3*Δ *rtt106*Δ, we next asked whether cells divided differently. We imaged cells over 16 h and tracked individual cells of each genotype from first septation, to the individual septation events of each daughter cell (Figure 5A). The first daughter to septate was labelled “D1” for daughter 1, and the other daughter was daughter 2, “D2”. We first examined proportions of dividing D1 and D2 cells in no treatment or with 12 mM HU or 10 µM CPT. We found no significant difference in numbers of D1 or D2 cells that divided in the absence of drug, although the *rad3*Δ *rtt106*Δ cells had higher D2 division (Figure 5B, dark grey bar, p=0.07) than *rad3*Δ alone. Ratios of D1/D2 show no difference in division.

**Figure 5.**
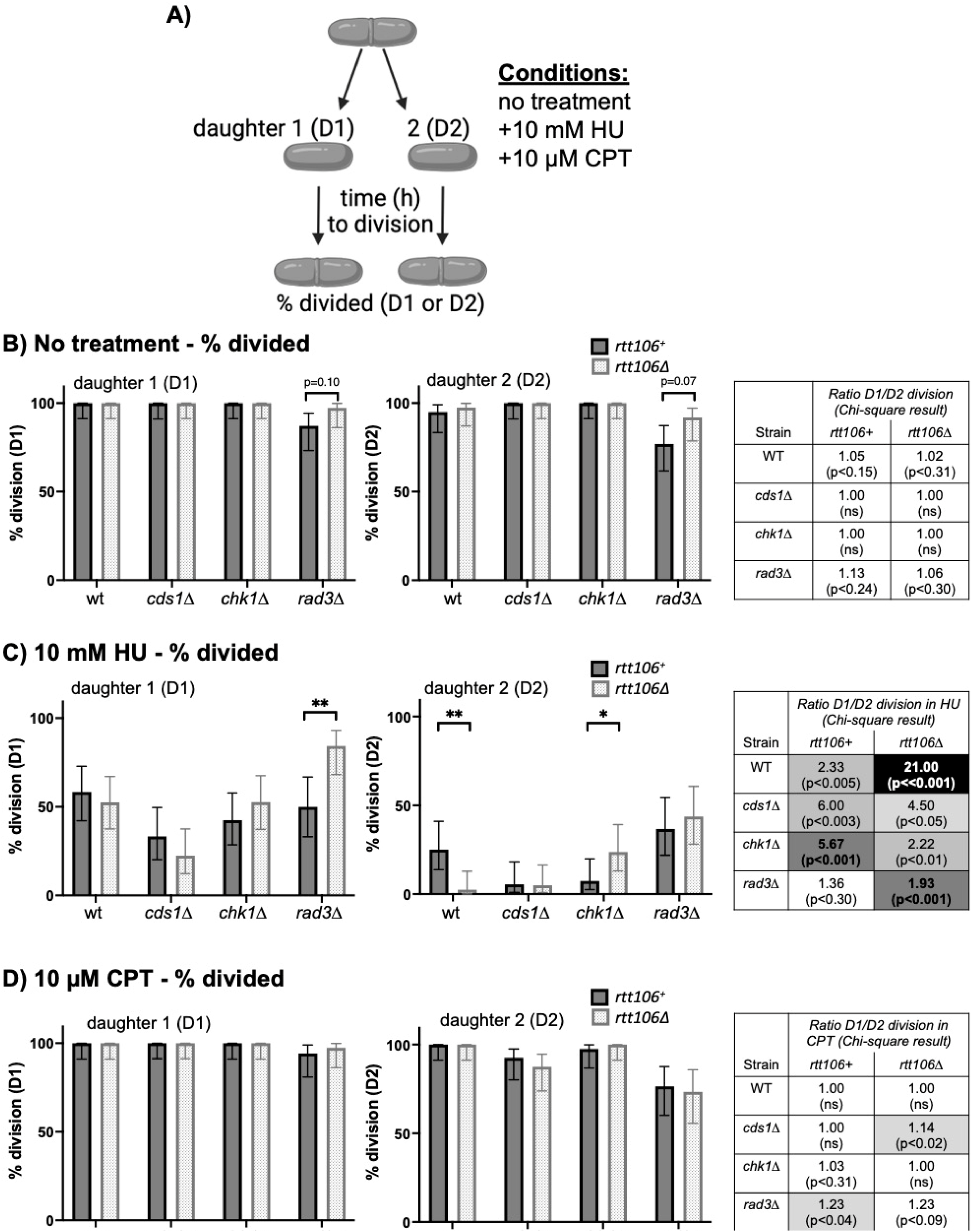
Rtt106 prevents cell division in the presence of DNA damage following replication stress. A) Generational division was assessed in no treatment, 10 mM hydroxyurea, and 10 μM camptothecin conditions. Single cells were tracked from an initial division to division of each daughter cell. The first daughter cell to divide is labeled daughter 1 (D1), with the other daughter cell D2. Timing of divisions is explored in Figure 6. Each data set in B, C, D is from 2 or 3 independent experimental replicates, with a minimum of 20 cell division sets counted per replicate. Daughters that septated by the end of 16h were scored relative to D1 or D2 status. B) In the no treatment group, there no significant difference between single and double mutants, nor between daughter 1 and daughter 2. The chart (right) compares the ratio of D1/D2 dividing cells for each strain. If the number of D1 and D2 divisions was the same, the ratio would be 1; more D1 divisions than D2 divisions produces a larger value. D1/D2 ratios are shown with a Chi-square p-value comparing dividing cells in each D1 or D2 population. C) In 10 mM HU, fewer cells divide generally, and *rad3*Δ *rtt106*Δ D1 cells are more likely to divide compared to all other genotypes. The *chk1*Δ *rtt106*Δ D2 cells are more likely to divide compared to *chk1*Δ alone. However, *rtt106*Δ D2 cells are less likely to divide in HU compared to wild type D2 cells, indicating tighter arrest in HU. D1/D2 ratios of each strain reinforce the observation of D1 or D2 division in HU. Darker grey chart cells indicate that the Chi-square comparison is less (white, p>0.05) to most (black, p<<0.001) significant. D) Cell division in CPT was observed and presented as in B and C. Although no percent division was significantly different between *rtt106+* and *rtt106*Δ in the charts, *cds1*Δ *rtt106*Δ D1 cells divided more than D2 in CPT.

Because HU affected growth, we tested cells in 12 mM HU. We found that wild type D1 cells were more likely to divide compared to D2 (Figure 5C, chart), a trend that was increased in *rtt106*Δ where the ratio of D1/D2 was 21. The loss of D2 division in *rtt106*Δ was significantly different from that of wild type in HU. The opposite was the case for *chk*Δ in HU; while D1 division was unaffected, *chk1*Δ *rtt106*Δ D2 cells were more likely to divide in HU than *chk1*Δ alone. D1 division was primarily affected in *rad3*Δ *rtt106*Δ, which had almost 100% D1 division compared to approximately 50% in *rad3*Δ. These data suggested that HU-replication stress causes DNA damage that affects primarily D2 cells. These cells were exposed before the initial septation event that formed both daughters. Because Rtt106 prevents some *rad3*Δ D1-division in HU, and *chk1*Δ D2-division, we conclude that HU generates DNA damage that promotes Rtt106-dependent arrest in D1 and D2 cells. Because *rad3*Δ*rtt106*Δ cells show a high proportion of DNA mis-segregation and “*cut*” morphologies like *rad3*Δ, we suggest that continued *rad3*Δ *rtt106*Δ D1 division occurs in a daughter that receives more DNA after the first disordered division.

Finally, we assessed CPT on D1 and D2 divisions. We found no significant differences that were dependent on Rtt106 status. However, we found that *cds1*Δ *rtt106*Δ D2-divisions are lower than the D1 divisions (Figure 5D, chart; p=0.02). Both *rad3*Δ strains show a similar ratio of D1/D2 division, indicating that the response of either daughter cell to CPT is similar in either *rad3*Δ or *rad3*Δ *rtt106*Δ cells.

Because RTT106 influences histone H3 levels, we hypothesized that deposition of H3 and CenpA may be altered at centromeres, which could alter cell division timing in the presence of replication stress or DNA damage. We tracked the time from parent septation to D1 and D2 septations and modeled the median division time for each strain with or without Rtt106 (Figure 6). We found that *rtt106*Δ cells had a 10 min longer median time to D1 division compared to wild type (Figure 6A). This indicated that Rtt106 might contribute to G2/M-checkpoint effects that prevent DNA damage in the subsequent daughter cell divisions. Agreeing with this prediction, the *rad3*Δ *rtt106*Δ D1-cells divided 50 min faster than *rad3*Δ alone. Similarly, *rad3*Δ *rtt106*Δ D1-cells in CPT also showed a faster median division time (2:10 compared to 2:35, p<0.05).

**Figure 6:**
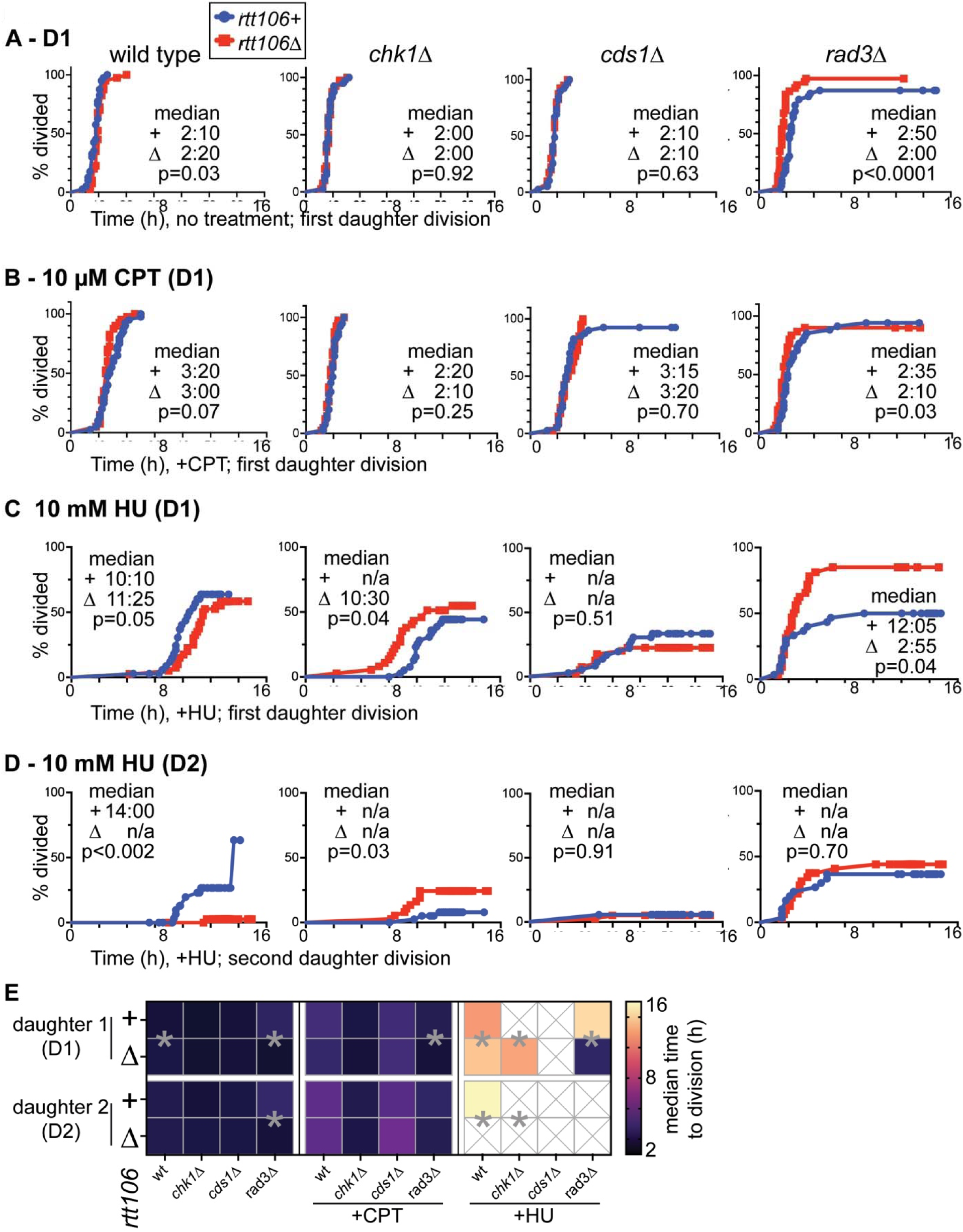
Rtt106 stops mitosis in the absence of the DNA repair checkpoint. Single fields of cells were observed in 16h movies following the scheme of Figure 5A. The time from initial septation to division of the first daughter (D1) or second daughter (D2) cell was recorded to show the % divided over time. Cultures of wild type, *chk1*Δ*, cds1*Δ*, or rad3*Δ, were observed after seeding, or in the presence of 10 µM camptothecin (CPT) or 10 mM hydroxyurea (HU). In all panels, blue is *rtt106+* and red is *rtt106*Δ for a given genotype. The median time to division was calculated from the curve and is presented in hh:mm. Cells that did not divide during the time of the movie were censored, and a given line may not attain 100% division. Cells that moved offscreen or were ambiguous were not used in the analysis. Curves were compared using the Log rank (Mantel-Cox) test of survival curve comparison to generate p values that indicate differences in the survival data timing and trends. A) D1 division was delayed in *rtt106*Δ cells without drug (far left, p=0.03). D1 division was unaffected in *cds1*Δ *and chk1*Δ cells (middle panels), but D1 median division time was 50 min faster in *rad3*Δ *rtt106*Δ cells (p=0.0001). B) CPT increased division time in D1 wild type and *cds1*Δ cells, while *chk1*Δ and *rad3*Δ D1 cells were slightly slower to divide. C, D) HU delayed D1 and D2 divisions in all cultures; curves with mostly censored cells that did not pass 50% divisions do not show a median time to division, shown as n/a. Wild type D1 and D2 divisions were faster than *rtt106*Δ. D1 and D2 median *chk1*Δ *rtt106*Δ division times were faster than *chk1*Δ alone. The *rad3*Δ *rtt106*Δ D1 cells were faster to divide and more likely to divide in HU compared to *rad3*Δ alone. E) Heat map of median time to division in D1 and D2 for each *rtt106+ and rtt106*Δ pair. Strains with a significantly different time to division have an asterix (*) between the *rtt106*+ and *rtt106*Δ. Conditions that did not generate a median time to division show “X” in a blank cell.

The biggest changes in division time relative to *rtt106*Δ occurred in HU. Time to division was longer in *rtt106*Δ compared to wild type for both D1 (1h 15 min faster) or D2 (Figure 6C, 6D). The *rtt106*Δ D2 cells failed to divide in HU, suggesting that HU-dependent DNA damage is more severe in D2 cells. No D1 or D2 changes were seen in *cds1*Δ compared to *cds1*Δ *rtt106*Δ in HU. However, *rad3*Δ *rtt106*Δ median division time was 9h faster than *rad3*Δ in HU D1 divisions, and ∼50% of *rad3*Δ D1 cells did not divide. D2 divisions were similar in *rad3*Δ regardless of Rtt106 (Figure 6D, 6E). This suggests that *rad3*Δ *rtt106*Δ D2 cells are similar to *rad3*Δ D2 cells treated with HU.Both D1 and D2 of *chk1*Δ *rtt106*Δ divided faster (D1) or more (D2) than *chk1*Δ in HU.

Because *cds1*Δ cells were unaffected in any condition with *rtt106*Δ, including in HU, we inferred that the DNA replication checkpoint is not affected by loss of Rtt106. Instead, the accelerated D1 division of *rad3*Δ *rtt106*Δ and *chk1*Δ *rtt106*Δ cells in a variety of conditions indicates that Rtt106 prevents precocious septation. Because *rtt106*Δ cells alone show delayed septation compared to wild type, we conclude that Rtt106 prevents DNA damage that causes cell cycle arrest in subsequent cell divisions.

## DISCUSSION

Histone chaperones maintain and stabilize DNA by contributing to chromatin structure folding (Gurard-Levin *et al*., 2014). We have found that Rtt106 histone chaperone promotes genome stability in fission yeast; in the absence of Rtt106, cells show increased mutation rate that is amplified by the loss of the DNA damage checkpoint (*rad3*Δ*, chk1*Δ). Because damaged DNA must be repaired to prevent mutations and chromosome changes (Kastan and Bartek, 2004), we propose that Rtt106 actively regulates histone H3 levels. We also found that *rtt106*Δ cells accumulate more histone H3 protein, a state that causes chromosome mis-segregation in other organisms. We found that Rtt106 prevents precocious DNA division in HU conditions that cause DNA damage. We conclude that Rtt106 regulates histone levels to promote genome stability.

Because of their role in chromatin structure and maintenance, histone and chaperone protein levels are increased in S-phase (Mei *et al*., 2017) and are regulated all other times (Franklin *et al*., 2021). Loss of core histone proteins can slow the cell cycle and decondense chromatin (Prado *et al*., 2017), effects that are connected to immune deficiency (Kim *et al*., 2021) and cellular senescence and aging (Dubey *et al*., 2024). Loss of histone variants such as centromeric CenpA can cause chromosome mis-segregation. Budding yeast histone chaperone RTT106 influences general transcription (Imbeault *et al*., 2008) and recruits chromatin modeling SWI/SNF and RSC complexes to histone genes during S-phase (Ferreira *et al*., 2011). Too much histone protein can also disrupt genome stability (Singh *et al*., 2010; Liang *et al*., 2012). Excess histones can bind to chromatin modifying enzymes and reduce their chromatin-specific functions (Singh *et al*., 2010). Direct links between excess histone and DNA damage signaling and processing have also been described. Drosophila Chk1 is competitively inhibited by excess H3 protein, which reduces *chk1*Δ function (Shindo and Amodeo, 2021). Budding yeast RAD53 binds to excess histones, and regulates their ubiquitination and degradation (Singh *et al*., 2009). Consequently, *rad53*Δ mutants cannot tolerate histone overexpression (Gunjan and Verreault, 2003; Singh *et al*., 2009). Although RAD53 is orthologous to fission *cds1*Δ, fission yeast *chk1*Δ is the primary DNA damage checkpoint kinase and better functional match to RAD53 for DSBs.

We predict that wild type and *rad3*Δ *rtt106*Δ cells higher mutation rates are caused by higher H3 levels in *rtt106*Δ. Excess histones impair homologous recombination repair by occluding repair sites and by non-specifically binding to repair proteins off of the chromatin (Liang *et al*., 2012). Precocious daughter cell division times in HU for *chk1*Δ and *rad3*Δ suggest that DNA damage is present but is potentially not detected with Rtt106 loss. Given the importance of H3 regulation to normal cells (Gunjan and Verreault, 2003; Singh *et al*., 2010; Liang *et al*., 2012) our data showing higher basal mutation rate in untreated wild type cells shows the importance of Rtt106 under regular cycling conditions. *S. pombe* expresses three histone H3 variants: H3.1, H3.2, and H3.3. Histone H3.3 deposition is independent of replication and nucleosome presence, and is associated with genomic instability through the activity of the DAXX histone chaperone (Voon and Wong, 2016; Molenaar *et al*., 2022). Since fission yeast dNTP metabolism, replication origins, RAD53 function, and centromere structures are different from budding yeast, future work will determine whether increased histone H3 in *rtt106*Δ similarly decreases DNA repair, or if centromere structure and DNA segregation are impacted with an altered spindle checkpoint.

## MATERIALS & METHODS

### Yeast strains and growth conditions

*rtt106*Δ single and double mutant *S. pombe* strains were generated from wild type, *cds1*Δ, *chk1*Δ*, rad3*Δ *S. pombe* strains, as described in Table 1. Growth and testing cells was performed as described in (Sabatinos and Forsburg, 2010; Petersen and Russell, 2016; Kianfard *et al*., 2022b). Cultures were grown overnight at 30°C with shaking at 200 rpm in glass flasks. Growth was in yeast extract with 225 mg/mL supplements (YES, with Ade, His, Leu, Ura, Lys) media unless otherwise described. Cells were maintained on YES plates with 0.8% agar. Before use, cultures were assessed for density using absorbance measurements at a wavelength of 600 nm (OD600), or by hemocytometer counting. Cultures were diluted in medium and recovered for 2 hours at 30°C with shaking if necessary to attain desired yeast growth stage, for example in HU acute survival.

### Spot test for drug sensitivities

Drug plates were made freshly in YES agar medium using filter sterilized drug stocks including: hydroxyurea (1M in water, BioShop, Canada); camptothecin (10 mM in DMSO; BioShop Canada); phleomycin (100 mg/mL in PBS, Invitrogen Zeocin). Cells were counted using a hemacytometer, and then diluted to normalize to 1 x 10^6^ cells/mL. Normalized cultures were serially diluted 1:5 in YES using a 96-well, and then pinned onto agar plates. Spots were grown at 30 for 3 – 5 days before scanning analysis.

### Western blot of histone levels

SDS-PAGE gels were made with 16% acrylamide and run in 1x Tris-Glycine buffer. Gels were electro-blotted onto PVDF membrane in transfer buffer (10% methanol, 10mM CAPS).

Blocking buffer and washes were performed using phosphate buffered saline with 0.1% Tween 20 (1x PBT). Membranes were blocked in 5% milk for 1 hour at room temperature. Antibody against histone H3 (rabbit, Cell Signal Technologies #97155) or PCNA (mouse, CST#25865) were added at 1:1000 in block for overnight incubation at 4°C. Membranes were washed 3 times in PBST, and then incubated in anti-rabbit or anti-mouse 1:5000 HRP-conjugated secondary antibody in block for 2 hours at 4°C. Membranes were washed 3 times in PBT and then imaged using Clarity ECL substrate (BioRad 1705060). Signal was detected on a BioRad Chemidoc.

### Canavanine forward mutation rate determination

Canavanine mutation rate was assess as described in (Karam and Sabatinos, 2025). Briefly, cells were grown to mid-exponential phase in PMG medium with supplements (PMG-S; His, Ade, Leu, Ura at 225 mg/L). Cultures were split, and half was treated with 12 mM HU for 4h at 30°C. Five millilitres of cultures were harvested and centrifuged to pellet cells. The pellets were resuspended in 50 µL of PMG-S. Twenty microlitres of resuspended culture was diluted 1:10,000 and plated onto YES agar plates to assess colony formation and cell number. The remaining amount was spotted onto PMG-S with 70 µg/mL canavanine, then dried and incubated 8 - 10 days at 30°C. Mutated colonies gain the ability to grow on canavanine. The number of *can*^+^ colonies was counted in each replicate and was compared to the viable cell number. Viable cell concentration was calculated from colony formation on YES titer dishes, grown 3 days at 30°C. The Lea and Coulson method of the median mutation rate per 10^7^ generations was calculated using FALCOR software (Hall *et al*., 2009). Mutation rates were compared using a 2-way ANOVA The Ma-Sandri-Sarkar Maximum Likelihood Estimator (MSS-MLE, (Hall *et al*., 2009)) was used to generate a best-estimate mutation rate per 10^7^ generations described in Table 2.

### Viability of rtt106Δ mutants in liquid cultures

Overnight YES cultures were used to inoculate 30 mL of EMM-HULA medium and then were grown overnight at 30°C with shaking. Cultures were checked for growth phase concentration, and 12 mM HU was added. Samples were removed at 0, 2, 4, and 6 hours, and diluted 1:10^3^ in YES. Diluted cultures were plated in duplicate onto 10 cm YES dishes and grown for 3 days at 30°C before colony counting. Viability was calculated for each strain relative to the 0h timepoint. Samples were fixed in 70% ethanol at each timepoint and stored at 4°C for imaging and flow cytometry (Kianfard *et al*., 2022a, 2022b).

Half-maximal inhibitory concentration (IC50) tests were performed in YES medium in 96 well plates, using the methods described (Chhipa *et al*., 2025). Data for each replicate were plotted in GraphPad Prism software, with a 3-parameter log curve. The area under the curve (AUC) for each replicate was calculated. Differences between *rtt106+* and *rtt106*Δ strains were compared using t-tests in GraphPad Prism. An aggregate IC50 value was plotted for each strain and drug, which considers the experimental variation between replicates.

Kinetic growth tests were performed in 96-well plates as described (Sanayhie and Sabatinos, 2025). Briefly, overnight cultures were resuspended at approximately 1 x 10^5^ cells/mL in wells with 0, 3, 6, or 12 mM HU. The plate was incubated at 30°C in a Synergy HTX (Agilent Canada) multimodal plate reader, with double orbital shaking for 16 hours. Optical density at 600 nm was measured every 15 min.

### Aniline blue/DAPI cell staining and imaging

Fixed *S. pombe* cells from acute viability timepoints were stained with aniline blue to detect septa, as described (Kianfard *et al*., 2022a). Briefly, fixed cells were rehydrated in PBS, incubated with 0.1% aniline blue for 10 minutes, and then centrifuged. Pelleted cells were dried onto glass coverslips, which were mounted onto slides with glycerol-based mounting media containing 0.1% DAPI. Slides were imaged on an Olympus microscope using a 60X oil immersion objective lens. DAPI and brightfield images were obtained for at least 3 separate fields of view in each replicate. Images were analyzed in FIJI software (Schindelin *et al*., 2012) to measure cell length. Images were counted for mono-, bi-, or anucleate, mitotic, and septated cells, in addition to abnormal DNA segregation patterns as described in (Kianfard *et al*., 2022a).

### Live cell imaging and division timing

Cultures were grown overnight to exponential phase in EMM+HULA medium. Optical density was used to normalize cultures to a similar starting value, approximately 1 x 10^6^ cells/mL. A 96-well plate was prepared with EMM+HULA medium alone, or with 10 mM hydroxyurea or 10 µM CPT. Drug concentration was calculated based on final volume in each well as described in (Sanayhie and Sabatinos, 2025). Culture was added to a final concentration of approximately 1 x 10^3^ cells/mL per well. The prepared plate was centrifuged at 1000 rpm for 1 minute before loading onto an environmental imaging chamber set to 32°C (OKO Labs, Italy). Cells were imaged using phase contrast on a Nikon Ti2 epifluorescent imaging system with 40x lens (Nikon Canada) every 10 minutes for 16 hours. Cultures were assessed in duplicate wells, and 3 to 4 experimental replicates were performed in total. Time lapse movies were analyzed using ImageJ to find stably septating cells. A minimum of 20 stable cells with defined endpoints were tracked in a minimum of 2 experimental replicates. Time of parental cell septation was noted, and then each daughter cell was tracked to determine time of its septation. The first daughter cell to septate was labelled “Daughter 1” (D1). The proportion of dividing D1 and D2 cells was calculated in GraphPad Prism, and differences in number of dividing cells (*rtt106+* versus *rtt106*Δ) were assessed using a Chi-squared test between each pair-wise comparison.

Time to division (D1 or D2) was calculated in Microsoft Excel, by subtracting the parental division time from the division time of D1 or D2. Times were imported into GraphPad Prism for survival analysis. Proportions of dividing cells were plotted over time using a Kaplan-Meyer curve. The median time to division was calculated from this curve. Pairs of *rtt106^+^* versus *rtt106*Δ curves were compared using log rank and Mantel-Hertzel tests of significance.

## ACKNOWLEDGMENTS

This work was funded by the Canadian Natural Science and Engineering Research Council (NSERC) to SAS (RGPIN-2015-04405) and JF (Discovery Grant Program). EK was supported by an Ontario Graduate Scholarship. This work was supported by TMU Faculty of Science NSERC Booster and Bridge support to SAS, and URO Funding to SSanayhie and NK.

## Abbreviations

HU: hydroxyurea
CPT: camptothecin
Phleo: phleomycin
D1/D2: daughter 1/ daughter 2
PMG: pombe minimal glutamate medium (with supplements, -S)
YES: yeast extract with supplements medium

